# Indirect genetic effects improve female extra-pair heritability estimates

**DOI:** 10.1101/2022.12.08.519585

**Authors:** Sarah Dobson, Jamie Dunning, Terry Burke, Heung Ying Janet Chik, Julia Schroeder

## Abstract

The question of why females engage in extra-pair behaviours is long-standing in evolutionary biology. One suggestion is that these behaviors are maintained through pleiotropic effects on male extra-pair behaviors and lifetime reproductive success (genes controlling extra-pair behaviours are shared between sexes, but only beneficial to one, in this case, males). However, for this to occur extra-pair behaviour must be heritable and positively genetically correlated between sexes. Although previous studies have suggested low heritability with no evidence for between-sex genetic correlations in extra-pair behaviours, indirect genetic effects (those derived from the behaviour of others, IGEs) from the social partner, the influence of the social partner’s genotype on the phenotype of an individual, have not been considered, despite the potential to uncover hidden heritability. Using data from a closed house sparrow population with a genetic pedigree spanning two decades, we tested the influence of IGEs on heritability and genetic correlation estimates of extra-pair behaviour. We found that the inclusion of IGEs improved model fit for both male and female extra-pair heritability. While IGEs did not change between-sex genetic correlations, we found a reduction in uncertainty in our estimates. Future studies should consider the effect of IGEs on the mechanisms of sex specific extra-pair behaviour.

## Background

The question of why females engage in extra-pair behaviour is long-standing in evolutionary biology (Griffith, Owens and Thuman 2002; Brouwer and Griffith 2019). Extra-pair reproductive behaviours are those which happen outside of an established social pair bond, from copulation to realized paternity of extra-pair offspring, and these behaviours are common in otherwise monogamous passerine groups (Griffith, Owens, and Thuman 2002; Cockburn 2006). Variation in extra-pair behaviour between and within species has been linked to phylogenetic variation (Brouwer and Griffith 2019),suggesting a genetic basis for the trait, but the mechanism of selection may vary between sexes. While the benefits of engaging in extra-pair behaviour is clear from the male perspective – siring more offspring without investing in costly parental care (Trivers 1972; Lebigre, Arcese, and Reid 2013; Raj Pant et al. 2022) – this is not the case for females, who instead replace within-pair offspring with extra pair offspring and risk associated costs (Trivers, 1972; Dixon et al. 1994; Valera, Hoi, and Krišti n 2003; Poiani and Wilks 2000; Matysioková and Remeš 2013; Schroeder et al. 2016; Albery et al. 2021). Despite appearing mal-adaptive, females actively seek extra-pair copulation (Lifjeld and Robertson 1992; Forstmeier 2007; Girndt et al. 2018), while the mechanisms driving the behaviour remain unresolved.

Adaptive hypotheses seek to explain female participation in extra-pair behaviours in the context of indirect benefits - those that benefit her offspring. However, evidence for such benefits is sparce (Akçay and Roughgarden 2007; Arct, Drobniak, and Cichoń 2015; Hsu et al. 2015; Grinkov et al. 2022). Indeed, studies even often suggest costs, both to extra-pair offspring (Schmoll et al. 2009; Sardell et al. 2012; Hsu et al. 2014), and to promiscuous females (Forstmeier 2007; Matysioková and Remeš 2013; Schroeder et al. 2016).

An alternative non-adaptive hypothesis posits that female extra-pair behaviours persist through intersexual pleiotropy (Halliday and Arnold 1987) - where two or more traits are controlled by associated sets of genes, present in both sexes. For intersexual pleiotropy to act as a driver, female extra pair behaviour would have to be linked to a male trait under strong positive selection, in this case, male extra-pair behaviours (Halliday and Arnold 1987; Reid and Wolak 2018). However testing this hypothesis is difficult because any study must demonstrate that male extra-pair behaviour is beneficial, that extra-pair behavior is heritable and positively genetically correlated between sexes (Forstmeier et al. 2014).

There is plenty support for male extra-pair behaviours contributing to male lifetime reproductive success (Albrecht et al. 2009; Webster et al. 2007; Lebigre, Arcese, and Reid 2013; Baldassarre and Webster 2013; Losdat, Arcese, and Reid 2015; Raj Pant et al. 2022), however, heritability estimates are low or not significant (Reid et al., 2011a; Reid et al., 2011b; Reid and Wolak, 2018; Grinkov et al., 2020). Further, those few existing empirical studies did not find evidence for between-sex correlation in extra-pair behaviour in captivity (Wang et al. 2020) nor in wild populations (Zietsch et al. 2015; Reid and Wolak 2018).

However, these studies did not include indirect genetic effects (IGEs), which represent part of the total additive genetic variance in a trait (Bijma 2010; Bailey, Marie-Orleach, and Moore 2018). IGEs are the influence of a conspecific’s genotype on the phenotype of an individual, often derived through the social environment (Maldonado-Chaparro et al. 2018) and which mediate trait expression (Wolf et al. 1998; Bijma 2010). Despite the potential of IGEs to uncover “hidden” genetic variation, most additive genetic variance, heritability and between sex genetic correlation estimates of extra-pair behaviours are calculated using only direct genetic effects, thus possibly under-estimating total genetic variation, (Kruuk and Wilson, 2018). Further, IGEs may facilitate selection even where direct heritability is low, with potential positive genetic covariance between the focal individual and their partner further accelerating trait evolution (Bijma 2010; Bailey, Marie-Orleach, and Moore 2018; Schroeder et al. 2019).

Here, we tested for heritability, then between-sex genetic correlation of extra-pair behaviours and how the inclusion of social partner IGEs influences both, in a closed and intensely monitored, wild island passerine population, with a genetic pedigree spanning two decades.

## Methods

### Study population

We have monitored the sedentary and closed population of house sparrow *Passer domesticus* (hereafter sparrow/s) breeding on Lundy Island in the Bristol Channel, UK (51°10’N, 4°40’W), since 1984 (see Ockendon, Griffith, and Burke 2009; Schroeder et al. 2012; Dunning et al. 2022). The sparrows on Lundy breed in nest boxes. All sparrows are marked with a unique sequence of three coloured leg rings and a BTO metal ring to allow subsequent identification of social pairs (for details see Nakagawa et al. 2007). Annually, nearly all present sparrows are detected by observation and capture, without bias (Simons et al. 2015). We collected tissue samples from nestlings at the natal site and from recaptured post-fledglings and used 22 microsatellite loci to allocate paternity (Dawson et al. 2012). We used these data to construct a genetic pedigree (Schroeder et al. 2015) spanning twenty years from 2000 – 2019, comprised of 8151 individuals, and where 931 individuals engaged in extra-pair events.

Female house sparrows are socially monogamous, but genetically promiscuous (Schroeder et al. 2016), and, on Lundy, our sparrows instigate a mean of 2.3 (sd 1.04) broods and lay a mean of 9.1 (sd 4.9) eggs annually (Westneat et al. 2014). Although previous studies have described extra-pair behavior (Ockendon, Griffith, and Burke 2009; Hsu et al. 2014, 2015, Schroeder et al. 2016), they find no evidence for adaptive benefits to females in this system. Extra-pair offspring were identified where the genetic sire differed from the social partner in the pedigree.

### Heritability and IGE models

We estimated within-sex heritability and between-sex genetic correlations of extra-pair behaviours using animal models. Animal models use a genetic pedigree within a mixed effect model to differentiate between environmental and genetic influences on a phenotypic trait (Wilson et al. 2010). We ran a series of animal models using MCMCglmm (Hadfield 2010) in R, v3.6.3 (R core team et al. 2022). We used a series of univariate and bivariate models with differing random effects to determine whether social partner IGEs improved within-sex extra-pair heritability estimates. We then used the same approach to test for genetic correlations between male and female extra-pair behavior (see Table 1 for model specifications). To improve confidence in our models we repeated each using multiple sets of priors, which confirmed results qualitatively (Table S1).

**Table 1.** Model specifications for extra pair behaviour (EPB) univariate models (1.1 - 4.2) and bivariate models (5-7). Responses and fixed effect specifications are universal across all models. Random effects are specified with different parameters for different models and include direct genetic effects (DGE), individual permanent environment (PE), social partner permanent environment (Partner PE) and social partner indirect genetic effects (Partner IGE). cov() denotes models where covariation between male and female parameters was allowed to take on any value. Idh() denotes were covariation between male and female parameters was fixed to 0.

| Model | Response | Random Effects |
| --- | --- | --- |
| 1.1 | Male EPB | male DGE + male PE |
| 1.2 | Female EPB | female DGE + female PE |
| 2.1 | Male EPB | male DGE + male PE + female partner PE |
| 2.2 | Female EPB | female DGE + female PE + male partner PE |
| 3.1 | Male EPB | male DGE + male PE + female partner PE + female partner IGE |
| 3.2 | Female EPB | female DGE + female PE + male partner PE + male partner IGE |
| 4.1 | Male EPB | cov(male DGE + female partner IGE) + male PE + female partner PE |
| 4.2 | Female EPB | cov(female DGE + male partner IGE) + female PE + male partner PE |
| 5 | Male EPB and<br>Female EPB | cov(male DGE + female DGE) + idh(male PE + female PE) |
| 6 | Male EPB and<br>Female EPB | cov(male DGE + female DGE) + idh(male PE + female PE) +<br>idh(female partner PE + male partner PE) |
| 7 | Male EPB and<br>Female EPB | cov(male DGE + female DGE) + idh(male PE + female PE) +<br>idh(female partner PE + male partner PE) + cov(female partner IGE +<br>male partner IGE) |

We measured male extra-pair behaviour using the number of extra-pair offspring within broods, but, to reflect the limited number of extra-offspring a female can produce, we measured her extra-pair behaviour as a threshold trait, recording presence/absence within broods. To reflect the increased likelihood of older males siring extra-pair offspring, we included age as a fixed effect across all models (Girndt et al. 2018). We did not include fine-scale environmental or social effects, which can potentially bias heritability estimates in closed systems (Germain et al. 2016; Grinkov et al. 2022).

In models measuring direct heritability we only included the focal individuals’ identity twice as random effects: the first to estimate the effect of the individual permanent environment and the other linked to a pedigree-based inverse relatedness matrix to estimate direct genetic effects (Table 1, Models 1.1 + 1.2). This allowed us to separate environmental and genetic causes of variance in extra-pair behaviour due to focal individual identity and to correctly estimate direct genetic effects (Kruuk and Hadfield, 2007). In models measuring total heritability we also included the identity of the social partner twice: one to estimate social partner permanent environment and the other linked to a pedigree-based relatedness matrix to estimate social partner indirect genetic effects. In univariate models we also modelled indirect genetic effects with covariation between direct and indirect genetic effects, but we could not do this in bivariate models due to MCMCglmm package constraints (Table 1). We constrained the covariance to zero? of all non-genetic effects in bivariate models because extra-pair behaviours are not expressed by the same individual (Table 1). Initial analysis included year as a random effect but as it accounted for little variation and did not change direct heritability and genetic correlation estimates across multiple models (Table S2) it was removed from further analysis.

We estimated heritability and genetic correlations using the equations listed in Table S3 on both the latent scale, using untransformed model outputs from non-gaussian models, and the phenotypic scale using the R package QGglmm (Villemereuil et al. 2016). All models were deemed to have converged when autocorrelation was less than 0.1, trace and density plots were unimodal and effective sample sizes for each effect were >1000 (Hadfield 2010). We assessed model fit using Deviance Information Criterion (DIC), where a decrease in value of more than 2 units indicates a better fit.

## Results

Our data contained 6774 offspring from 1923 broods between 793 unique breeding pairs. Of those offspring, 19% (1287) were extra-pair from 41% of broods (325). Changing social partners was common, with 47% of individuals having more than 1 social partner across their lifetime (Schroeder et al. 2016). Using the parameters detailed in the methods, we identified 3233 extra-pair events, between 776 individuals and over 18 years (1721 observations from 410 females and 1512 observations from 366 males) for use in our models.

Model outputs did not differ between models using different priors (Table 2, S4-S6). Direct additive genetic variance for male and male and female extra-pair behaviour were close to zero MTable 2). However, the addition of social IGEs increased model fit for male and female extra-pair behaviour (Table 2) and increased the total additive genetic variance available for both by 372% and 37. 5% respectively (Figure 1; Table 2),

**Table 2:** DIC, phenotypic scale direct heritability (h^2^_phen_), latent scale direct heritability (h^2^_lat_), phenotypic scale total heritability (t^2^_phen_), latent scale total heritability (t^2^_lat_), direct additive genetic variance (V_Adge_), indirect genetic effect variance (V_Aige)_, permanent environment (Vpe), social permanent environment (Vse) and covariation between direct and indirect genetic effects (cov(V_Adge,_ V_Aige)_) from the posterior means of extra pair behaviour (EPB) univariate models using prior 1. Heritability and variance estimate where CIs were ≥0.001 are highlighted in bold. Residual effect, intercept and fixed effect estimates can be found in Table S5.

| Models | Response | Parameters | DIC | $h^2_{\text{phen}}$<br>$/t^2_{\text{phen}}$ | $h^2_{\text{lat}}$<br>$/t^2_{\text{lat}}$ | $V_{\text{Adge}}$ | $V_{\text{Aige}}$ | $V_{\text{pe}}$ | $V_{\text{se}}$ | Cov<br>($V_{\text{Adge}}$,<br>$V_{\text{Aige}}$) |
| --- | --- | --- | --- | --- | --- | --- | --- | --- | --- | --- |
| 1.1 | **Male EPB** | **DGE +**<br>**PE** | 18<br>89 | 0.004<br>(<0.0<br>01 –<br>0.016) | 0.024<br>(<0.0<br>01 –<br>0.1) | 0.058<br>(<0.0<br>01 –<br>0.234) |  | **0.675**<br>**(0.323**<br>**–1.04)** |  |  |
| 1.2 | **Female**<br>**EPB** | **DGE +**<br>**PE** | 22<br>80 | 0.012<br>(<0.0<br>01–<br>0.042) | 0.02<br>(<0.0<br>01–<br>0.07) | 0.024<br>(<0.0<br>01 –<br>0.082) |  | **0.171**<br>**(0.065**<br>**–**<br>**0.28)** |  |  |
| 2.1 | **Male EPB** | **DGE +**<br>**PE +**<br>**Partner**<br>**PE** | 18<br>83 | 0.006<br>(<0.0<br>01<br>–<br>0.02) | 0.04<br>(<0.0<br>01 –<br>0.13) | 0.098<br>(<0.0<br>01–<br>0.324) |  | 0.363<br>(<0.0<br>01 –<br>0.71) | **0.614**<br>**(0.243**<br>**–**<br>**0.983)** |  |
| 2.2 | **Female**<br>**EPB** | **DGE +**<br>**PE +**<br>**Partner**<br>**PE** | 22<br>76 | 0.014<br>(<0.0<br>01 –<br>0.044) | 0.023<br>(<0.0<br>01–<br>0.07) | 0.029<br>(<0.0<br>01 –<br>0.087) |  | **0.15**<br>**(0.031**<br>**–0.27)** | 0.053<br>(<0.0<br>01 –<br>0.133) |  |
| 3.1 | **Male EPB** | **DGE +**<br>**PE**<br>**+**<br>**Partner**<br>**PE +**<br>**Partner**<br>**IGE** | 18<br>80 | 0.028<br>(<0.0<br>01 –<br>0.056) | **0.14**<br>**(0.016**<br>**–**<br>**0.29)** | 0.102<br>(<0.0<br>01 –<br>0.344) | 0.38<br>(<0.0<br>01–<br>0.82) | 0.355<br>(<0.0<br>01 –<br>0.7) | 0.275<br>(<0.0<br>01–<br>0.702<br>2) |  |
| 3.2 | **Female**<br>**EPB** | **DGE +**<br>**PE**<br>**+**<br>**Partner**<br>**PE +** | 22<br>77 | 0.022<br>(<0.0<br>01–<br>0.056) | 0.035<br>(<0.0<br>01–<br>0.09) | 0.032<br>(<0.0<br>01–<br>0.094) | 0.01<br>2<br>(<0.0<br>01–<br>0.04) | **0.149**<br>**(0.032**<br>**–0.27)** | 0.048<br>(<0.0<br>01 –<br>0.123) |  |
|  |  | **Partner**<br>**IGE** |  |  |  |  |  |  |  |  |
| 4.1 | **Male EPB** | **cov(Partner IGE and DGE) + PE + Partner PE** | 1881 | 0.017<br>(<0.001 – 0.040) | **0.114**<br>**(0.001 – 0.28)** | 0.1521<br>(<0.001 – 0.408) | 0.4343<br>(<0.001 – 0.84) | 0.284<br>(<0.001 – 0.67) | 0.2436<br>(<0.001 – 0.638) | -0.149<br>(-0.423 – 0.044) |
| 4.2 | **Female EPB** | **cov(Partner IGE and DGE) + PE + Partner PE** | 2277 | 0.019<br>(<0.001 – 0.052) | 0.029<br>(<0.001 – 0.08) | 0.029<br>(<0.001 – 0.09) | 0.012<br>(<0.001 – 0.04) | **0.1487**<br>**(0.037 – 0.26)** | 0.051<br>(<0.001 – 0.129) | -0.0017<br>(-0.025 – 0.019) |

**Figure 1):**
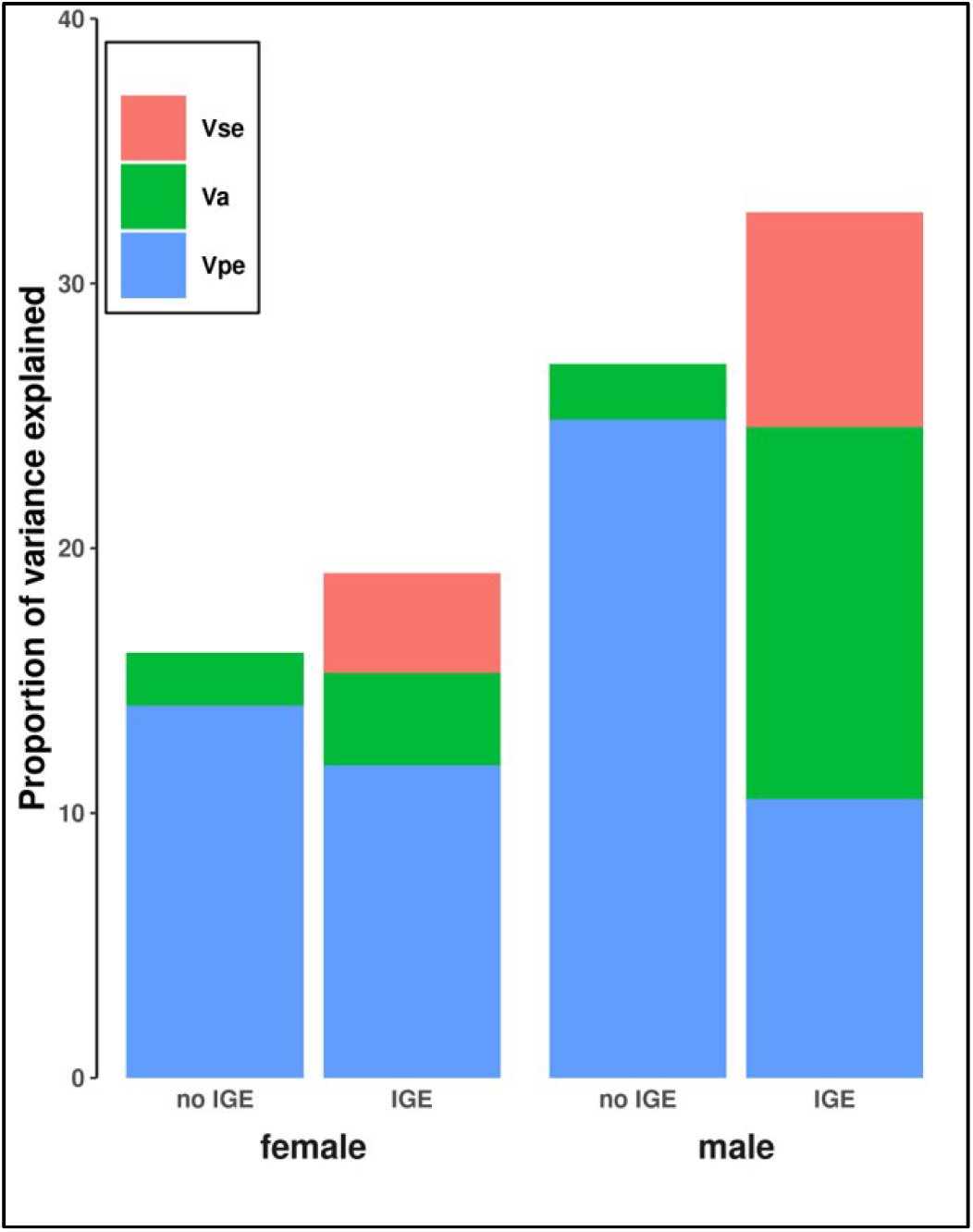
The proportion of the variance explained in male and female extra-pair behaviour in models without IGEs vs with IGEs on latent scales. Proportions include social partner permanent environment variance (Vse), additive genetic variance (Va) and direct permanent environment variance (Vpe). In bars without IGEs (those representing direct genetic effects) Va is calculated using models 1.1 and 1.2. In bars with IGEs (those that include total genetic variance) Va is calculated using models 3.1 and 3.2. Va is calculated using outputs from model 1 and model 3. Fixed effects accounted for in models, but not shown here.

Male and female extra-pair behaviours were mostly explained by social IGEs and individual permanent environment effects in the best fitting models(Table 2), accounting for 12% and 11% of total phenotypic variance respectively (Figure 1; Table 2). All models detected high levels of residual variation for both male and female extra-pair behaviour (Table S5).

### Male and Female Heritability

Both latent and phenotypic scale direct heritability estimates without social partner IGEs were close to zero for male and female extra-pair behaviour (Table 2). Social Partner IGEs slightly increased female total heritability estimates, but increased male total heritability substantially, however CIs were still close to zero for both males and females (Table 2). Covariation between direct and indirect genetic effects for male and female extra-pair behaviors were both negative but overlapped 0, reducing total heritability estimates (Table 2).

### Genetic Correlation Estimates between Male and Female extra-pair behaviour

Genetic correlations between male and female extra-pair behaviour estimated from direct genetic effects were positive, but CIs greatly overlapped zero (Table 3). The addition of social IGEs produced slightly negative correlations and CIs still overlapped zero but reduced the uncertainty in those estimates (Table 3). Heritability estimates did not differ between different priors, nor between univariate and bivariate models (Tables 2,3, S4-S6).

**Table 3:** DIC, phenotypic scale direct heritability (h^2^_phen)_, latent scale direct heritability (h^2^_lat_), phenotypic scale total heritability (t^2^_phen_), latent scale total heritability (t^2^_lat_), phenotypic scale genetic correlations (Vg_phen_) and latent genetic correlations (Vg_lat_) from the posterior means estimates of extra pair behaviour (EPB) bivariate models using prior 1. Heritability estimates where CIs were ≥0.001 are highlighted in bold. Intercept, residual and fixed effect estimations can be found in Table S5

| **Mo** | **Response** | **Parameters** | **DIC** | $h^2_{\text{phen}}/t^2_{\text{phen}}$ | $h^2_{\text{lat}}/t^2_{\text{lat}}$ | $V_{g\text{phen}}$ | $V_{g\text{lat}}$ |
| --- | --- | --- | --- | --- | --- | --- | --- |
| **dels** | (s) |  |  |  |  |  |  |
| 5 | Male | **cov(male** | 4171 | Male: 0.003 | Male: 0.02 | 0.09 | 0.089 |
|  | EPB and | **DGE +** |  | (<0.001 – | (<0.001– | (-0.84) – | (-0.84) – |
|  | female | **female** |  | 0.011) | 0.07) | 0.94 | 0.943 |
|  | EPB | **DGE) +** |  |  |  |  |  |
|  |  | **idh(male** |  | Female: |  |  |  |
|  |  | **PE +** |  | 0.007 | Female: |  |  |
|  |  | **female PE)** |  | (<0.001 – | 0.021 |  |  |
|  |  |  |  | 0.024) | (<0.001 – |  |  |
|  |  |  |  |  | 0.074) |  |  |
| 6 | Male | **cov(male** | 4160 | Male: 0.006 | Male: 0.039 | 0.045 | 0.045 |
|  | EPB and | **DGE +** |  | (<0.001 – | (<0.001 – | (-0.81) – | (-0.81) –0.97 |
|  | female | **female** |  | 0.02) | 0.125) | 0.97 |  |
|  | EPB | **DGE) +** |  |  |  |  |  |
|  |  | **idh(male** |  | Female: | Female: |  |  |
|  |  | **PE +** |  | 0.008 | 0.024 |  |  |
|  |  | **female PE)** |  | (<0.001 – | (<0.001 – |  |  |
|  |  | **+** |  | 0.026) | 0.073) |  |  |
|  |  | **idh(female** |  |  |  |  |  |
|  |  | **partner PE** |  |  |  |  |  |
|  |  | **+ male** |  |  |  |  |  |
|  |  | **partner** |  |  |  |  |  |
|  |  | **PE)** |  |  |  |  |  |
| 7 | Male | **cov(male** | 4157 | **Male: 0.032** | **Male: 0.138** | 0.066 | 0.066 |
|  | EPB and | **DGE +** |  | **(0.004 –** | **(0.02 –** | (-0.67 – | (-0.68 – |
|  | female | **female** |  | **0.064)** | **0.24)** | 0.55) | 0.55) |
|  | EPB | **DGE) +** |  |  |  |  |  |
|  |  | **idh(male** |  | Female: | Female: |  |  |
|  |  | **PE +** |  | 0.021 | 0.032 |  |  |
|  |  | **female PE)** |  | (<0.001 – | (<0.001 – |  |  |
|  |  | **+** |  | 0.057) | 0.085) |  |  |
|  |  | **idh(female** |  |  |  |  |  |
|  |  | **partner PE** |  |  |  |  |  |
|  |  | **+ male** |  |  |  |  |  |
|  |  | **partner** |  |  |  |  |  |
|  |  | **PE) +** |  |  |  |  |  |
|  |  | **cov(female** |  |  |  |  |  |
|  |  | **partner** |  |  |  |  |  |
|  |  | **IGE + male** |  |  |  |  |  |
|  |  | **partner** |  |  |  |  |  |
|  |  | **IGE)** |  |  |  |  |  |

## Discussion

Our results suggest that additive genetic variance and heritability for extra-pair behaviours are improved with the inclusion of social partner IGEs in both sexes, but particularly in males, where total genetic variation reached 14%. However, the addition of IGEs also increased uncertainty in variance. Although that uncertainty may be linked to increased model complexity, simulation studies suggest (Bijma *et al*., 2013) that 793 breeding pairs are sufficient to consider our estimates reliable. Our results support those of other systems, which find marginal heritability in extra-pair behaviours with high residual variation, implying that male and female extra-pair behaviours may be very flexible traits (Forstmeier *et al*., 2011, Reid *et al*., 2011a; Reid *et al*., 2011b; Zietsch *et al*., 2015; Grinkov *et al*., 2020; Wang *et al*., 2020; Beck *et al*., 2020).

Extra-pair reproduction may be driven by genes that control copulation and solicitation (Dixon *et al*., 1994; Matysioková and Remeš, 2013; Schroeder *et al*., 2016) but also by the behaviour of neighbors (Beck *et al*., 2020; Beck *et al*., 2021). However, not all copulations result in extra-pair offspring, due to mate guarding (Forstmeier *et al*., 2011) and post-copulatory processes (Forstmeier and Ellegren, 2010; Girndt *et al*., 2019). Therefore, while extra-pair copulations are probably common (Fossøy, Johnsen, and Lifjeld, 2006), copulations are unaccounted for in estimates (Girndt *et al*., 2019). Consequently, variance could be negatively biased against copulation behaviours (Forstmeier *et al*., 2014; Beck *et al*., 2020), affecting heritability. This has been demonstrated in captive zebra finches *Taeniopygia* sp., where heritability of copulation behaviours is substantial (Forstmeier *et al*., 2011; Wang *et al*., 2020), but equivalent data is extremely difficult to collect in the wild (Beck *et al*., 2020). Despite this, because genes are only passed on to the next generation through recruitment of extra-pair offspring, measurements in the wild have evolutionary relevance over copulations in captivity. Captive systems may also miss population scale processes, like assortative mating, which may skew heritability estimates for both extra-pair copulations and successful reproduction (Reid and Wolak, 2018; Wang *et al*., 2020).

Between-sex genetic correlations are notoriously difficult to estimate in wild populations due to the requirement of large sample sizes (Lynch, 1999; Bonduriansky and Chenoweth 2009; Morrisey *et al*., 2014), resulting in uncertainty in existing estimates (this study; Forstmeier *et al*., 2011; Reid and Wolak, 2018). Estimating between-sex genetic correlations between direct measures of male and female EPC may yield more precise estimates (Forstemeier *et al*., 2014). The inclusion of social IGE effects in male and female EPC measurements may reduce uncertainty even further. However, a high degree of uncertainty may also possibly hide existing weak positive or negative genetic correlations between female and male extra-pair behaviours (Reid and Wolak, 2018). Future work could consider simulations to aid answering these questions (Reid and Wolak, 2018), while wild sample sizes increase with subsequent generations.

Even if between-sex genetic correlations existed, pleiotropic effects on male lifetime reproductive success mean that they probably would not be maintained. For example, in pursuit of extra-pair copulations males may reduce mate guarding, resulting in fewer within pair offspring (Møller and Birkhead, 1993; Møller and Ninni, 1998; Harts and Kokko, 2013; Reid and Wolak, 2018). However, genes associated with siring extra- and within-pair offspring are correlated in some socially monogamous populations (Reid and Losdat, 2014; Reid and Wolak, 2018), suggesting that the mating strategies are plastic and may not change total male lifetime reproductive success. This combined, with low extra-pair offspring fitness (Hsu *et al*., 2015) could result in detrimental male lifetime reproductive success. It is also possible that female extra-pair behaviours persist through additional pleiotropic effects that benefit female fecundity (Forstemier *et al*., 2014). Positive genetic correlations between female solicitation behaviour and female fecundity have been described in captive populations (Wang *et al*., 2020), but as yet, empirical evidence from wild systems is lacking.

We used DIC to compare model fit, which we determined was a good measure of relative change between random effects. However, DIC may sometimes be unreliable where fixed effects or model families vary (Hadfield Pers. Comm.). Although this isn’t the case for our models, future work might also consider alternative approaches in determining model fit with varying random effects.

Our study explored the genetic basis and role of the social partner on male and female extra-pair behaviours, to better understand how such behaviours are maintained in socially monogamous populations. While we found no support for the notion that female extra-pair behaviours are maintained in socially monogamous populations through antagonistic intersexual pleiotropy. However, we suggest that social partner IGEs can uncover hidden genetic variation, especially for males. Social partner IGEs contributed substantially more to total male extra-pair heritability, accounting for more total additive genetic variance than male direct genetic effects and explained the largest proportion of phenotypic variation. We demonstrate the importance of IGE inclusion in quantitative models exploring aspects of animal behaviour. Future studies into why female extra-pair behaviours persist in socially monogamous populations should consider the intrasexual antagonistic pleiotropy hypothesis and, where sample size allows, should include social partner IGEs when estimating heritabilities.

## Acknowledgments

We thank the Lundy Landmark Trust and the Lundy Field Society. We also thank Issey Whinney. This research was supported by the NERC Quantitative Methods in Ecology and Evolution (QMEE) CDT, grant number NE/P012345/1 (JD), a fellowship from the Volkswagen Foundation (JS), a grant from the German Research Foundation: Deutsche Forschungsgemeinschaft (JS), CIG PCIG12-GA-2012-333096 from the European Research Council (JS)

## Code and data availability

A copy of the data and code used in this project can be found at https://github.com/Sazz01/CMEECourseWork/tree/master/research_project.

