## Supplementary material for "Indirect genetic effects improve female extra-pair heritability estimates": supp files

**Table S1: Structure of priors, number of iterations, thin and burnin specified for each MCMCglmm model. G and R represent G- structure and R- structure respectively**

| Model | Prior 1 | | Prior 2 | | nitt | thin | burnin |
| --- | --- | --- | --- | --- | --- | --- | --- |
|  | G | R | G | R |  |  |  |
| 1.1 | V = 1, nu = 1, alpha.mu = 0, alpha.V = 1 | V = 1, nu = 1 | V = 0.01, nu = 1, alpha.mu = 0, alpha.V = 1000 | V = 1, nu = 1 | 205000 | 200 | 5000 |
| 1.2 | V = 1, nu = 1, alpha.mu = 0, alpha.V = 1 | V = 1, fix = 1 | V = 0.01, nu = 1, alpha.mu = 0, alpha.V = 1000 | V = 1, fix = 1 | 310000 | 200 | 10000 |
| 2.1 | V = 1, nu = 1, alpha.mu = 0, alpha.V = 1 | V = 1, nu = 1 | V = 0.01, nu = 1, alpha.mu = 0, alpha.V = 1000 | V = 1, nu = 1 | 505000 | 500 | 5000 |
| 2.2 | V = 1, nu = 1, alpha.mu = 0, alpha.V = 1 | V = 1, fix = 1 | V = 0.01, nu = 1, alpha.mu = 0, alpha.V = 1000 | V = 1, fix = 1 | 1055000 | 1000 | 5000 |
| 3.1 | V = 1, nu = 1, alpha.mu = 0, alpha.V = 1 | V = 1, nu = 1 | V = 0.01, nu = 1, alpha.mu = 0, alpha.V = 1000 | V = 1, nu = 1 | 1055000 | 1000 | 5000 |
| 3.2 | V = 1, nu = 1, alpha.mu = 0, alpha.V = 1 | V = 1, fix = 1 | V = 0.01, nu = 1, alpha.mu = 0, alpha.V = 1000 | V = 1, fix = 1 | 2555000 | 2000 | 5000 |
| 4.1 | V = 1, nu = 1, alpha.mu = 0, alpha.V = 1 | V = 1, nu = 1 | V = 0.01, nu = 1, alpha.mu = 0, alpha.V = 1000 | V = 1, nu = 1 | 6055000 | 3000 | 5000 |
| 4.2 | V = 1, nu = 1, alpha.mu = 0, alpha.V = 1 | V = 1, fix = 1 | V = 0.01, nu = 1, alpha.mu = 0, alpha.V = 1000 | V = 1, fix = 1 | 6055000 | 3000 | 5000 |
| 5 | V=diag(2), nu= 2, alpha.mu = c(0, 0), alpha.V = diag(2) | V = diag(2), nu = 1.002, fix = 2 | V = diag(2)*0.02, nu = 3, alpha.mu = c(0, 0), alpha.V = diag(2)*1000 | V = diag(2), nu = 1.002, fix = 2 | 755000 | 500 | 5000 |
| 6 | V=diag(2), nu= 2, alpha.mu = c(0, 0), alpha.V = diag(2) | V = diag(2), nu = 1.002, fix = 2 | V = diag(2)*0.02, nu = 3, alpha.mu = c(0, 0), alpha.V = diag(2)*1000 | V = diag(2), nu = 1.002, fix = 2 | 505000 | 500 | 5000 |
| 7 | V=diag(2), nu= 2, alpha.mu = c(0, 0), alpha.V = diag(2) | V = diag(2), nu = 1.002, fix = 2 | V = diag(2)*0.02, nu = 3, alpha.mu = c(0, 0), alpha.V = diag(2)*1000 | V = diag(2), nu = 1.002, fix = 2 | 2555000 | 2000 | 5000 |

**Table S2: Comparison of phenotypic scale heritability estimates (H^2^) and genetic correlation estimates (Vg_lat_)) + 95% confidence intervals between univariate models with year as a random effect vs without year as a random effect**

| **Model** | **With Year** | **No Year** |
| --- | --- | --- |
| 1.1 | H^2^: 0.004 (<0.001 –0.016) | H^2^: 0.004 **(<**0.001 – 0.016) |
| 1.2 | H^2^: 0.007(<0.001 – 0.022) | H^2^: 0.007 (<0.001 – 0.023) |
| 2.1 | H^2^: 0.007 (<0.001 – 0.02) | H^2^: 0.006 (<0.001 – 0.02) |
| 2.2 | H^2^: 0.009 (<0.001 – 0.029) | H^2^: 0.008 (<0.001 – 0.025) |
| 5 | Male H^2^: 0.004 (<0.001 – 0.015)  Female H^2^: 0.007 (<0.001 – 0.025)  Vg: 0.04(- 0.8 - 0.8) | Male H^2^: 0.003 (<0.001 – 0.011)  Female H^2^: 0.007 (<0.001 – 0.024)  Vg: 0.09 (-0.84 - 0.94) |
| 6 | Male H^2^: 0.004(<0.001 – 0.013)  Female H^2^: 0.008(<0.001 – 0.024)  Vg_lat_: 0.01 (-0.7 – 0.9) | Male H^2^**:** 0.006 (<0.01 – 0.02)  Female H^2^: 0.008(<0.001 – 0.026)  Vg_lat_**:** 0.045 (-0.81 – 097) |

**Table S3: list of equations used to calculate heritability and genetic correlation estimates on latent and phenotypic scales adapted from Bijma (2010), De Villemereuil *et al.* (2018), and Schroeder *et al.* (2019). Where V_p_ is the total phenotypic variation, V_random_ is the combined variance of the random effects, V_fixed_ is the combined variance of the fixed effects, V_(Xb)_ is the variance of the linear predictor of the model, V_Adge_ is the direct genetic effects, V_Aige_ is the indirect genetic effects from the social partner, V_Athv_ is the total heritable variation, Cov_(VAdge * VAige)_ is the covariance between the direct and indirect genetic effects, V_Af_ and V_Am_ are the variances of the female and male genetic effects and Cov_(VAf * VAm)_ is the covariance between female and male genetic effects. Latent estimates were calculated as shown below. Estimates on the phenotypic scale were calculated using back transformed V_A_, V_P_, V_IGE_, Cov_(fDE * fDE)_, V_fA_ and V_mA_ estimates using QGglmm.**

| **Estimate** | **Equation** | **Applicable models** |
| --- | --- | --- |
| Latent scale fixed effect variance (V _fixed_) | $V_{fixed}=V_{\left( Xb \right)}$ | All models |
| Total phenotypic variation (V_p_) | $V_{P}=V_{fixed}+ V_{random}$ | All models |
| Total Heritable Variation (V_Athv)_ | $V_{Athv}=V_{Adge}+ V_{Aige}$ | 3.1, 3.2, 4.1, 4.2, 7 |
| Direct heritability (H^2^) | $H^{2}=\frac{V_{Adge}}{V_{P}}$ | 1.1, 1.2, 2.1, 2.2, 5, 6 |
| Heritability (H^2^) | $H^{2}=\frac{V_{Athv}}{V_{P}}$ | 3.1, 3.2, 7 |
| Heritability (H^2^) | $H^{2}=\frac{V_{Athv}+ 2Cov_{\left( V_{Adge}, V_{Aige} \right)}}{V_{P}}$ | 4.1, 4.2 |
| Genetic Correlation (V_g)_ | $V_{g}=\frac{Cov\left( V_{fAdge}, V_{mAdge} \right)}{\sqrt{\left( V_{fAdge}\cdot V_{mAdge} \right)}}$ | 5, 6 |
| Genetic Correlation (V_g_) | $V_{g}=\frac{Cov\left( V_{fAthv}, V_{mAthv} \right)}{\sqrt{\left( V_{fAthv}\cdot V_{mAthv} \right)}}$ | 7 |

**Table S4: DIC, Phenotypic scale heritability (H^2^_phen_), latent scale heritability (H^2^_lat_), direct additive genetic variance (V_Adge_), indirect genetic effect variance (V_Aige)_, permanent environment (Vpe), social permanent environment (Vse) and covariation between direct and indirect genetic effects (cov(V_Adge,_ V_Aige)_) from the posterior means of univariate models using Prior 2. Heritability and variance estimates where CIs were ≥0.001 are highlighted in bold. Residual effect, intercept and fixed effect estimates can be found in Table S5.**

| **Model** | **Response(s)** | **Parameters** | **DIC** | **H^2^ _phen_** | **H^2 l^at** | **Vadge** | **Vaige** | **Vpe** | **Vse** | **Cov(Vadge, Vaige)** |
| --- | --- | --- | --- | --- | --- | --- | --- | --- | --- | --- |
| **1.1**  **1.2** | **Male EPR**  **Female EPR** | **DGE + PE**  **DGE + PE** | 1888  2279 | 0.004 (<0.001 -0.014)  0.013  (<0.001 -0.041) | 0.022 (<0.001 - 0.08)  0.02  (<0.001-0.065) | 0.061 (<0.001 - 0.218)  0.024  (<0.001 - 0.079) | --  -- | **0.699 (0.365 - 1.123)**  **0.17**  **(0.067 - 0.286)** | --  -- | --  -- |
| **2.1**  **2.2** | **Male EPR**  **Female EPR** | **DGE + PE + Partner PE**  **DGE + PE + Partner PE** | 1882  2276 | 0.007  (<0.001 -0.021)  0.015  (<0.001 -0.047) | 0.034 (<0.001 -0.114)  0.02 (<0.001 -0.065) | 0.106 (<0.001 - 0.36)  0.03  (<0.001- 0.095) | --  -- | 0.389  (<0.001-0.736)  **0.153**  **(0.033 - 0.272)** | **0.625 (0.26 - 1.016)**  0.054  (<0.001 - 0.136) | --  -- |
| **3.1**  **3.2** | **Male EPR**  **Female EPR** | **DGE + PE**  **+ Partner PE + Partner IGE**  **DGE + PE**  **+ Partner PE + Partner IGE** | 1879  2277 | 0.029  (<0.001 -0.056)  0.021  (<0.001 -0.059) | **0.147**  **(0.001 - 0.29)**  0.034 (<0.001 -0.095) | 0.111 (<0.001 - 0.38)  0.03  (<0.001 -0.098) | 0.404  (<0.001-0.89)  0.0125  (<0.001 -0.045) | 0.367 (<0.001 -0.724)  **0.151**  **(0.035 - 0.285)** | 0.282 (<0.001 - 0.706)  0.05  (<0.001 - 0.126) | --  -- |
| **4.1**  **4.2** | **Male EPR**  **Female EPR** | **cov(Partner IGE and DGE) + PE + Partner PE**  **cov(Partner IGE and DGE) + PE + Partner PE** | 1880  2278 | 0.02 (<0.001 -0.043)  0.02  (<0.001 -0.057) | **0.134**  **(0.001 - 0.29)**  0.032  (<0.001 -0.092) | 0.143 (<0.001 - 0.42)  0.031  (<0.001 - 0.096) | 0.428  (<0.001 -0.086)  0.012  (<0.001 -0.046) | 0.315  (<0.001-0.675)  **0.15**  **(0.044 - 0.277)** | 0.265 (<0.001 - 0.679)  0.05  (<0.001 - 0.129) | -0.114 (-0.354 - 0.08)  -0.0015  (-0.027 -0.018) |

**Table S5: Intercept, Fixed effect and residual effect estimates for univariate and bivariate MCMCglmm models.**

| **model** | **Intercept/ Age** | | **Residual Effects** | |
| --- | --- | --- | --- | --- |
|  | **Prior 1** | **Prior 2** | **Prior 1** | **Prior 2** |
| **1.1** | Intercept: -3.046(-3.46 - -2.68)  pMCMC: <0.001 ***  Age: 0.312 (0.21 - 0.423)  pMCMC: <0.001 *** | Intercept: -3.049(-3.46 - -2.655)  pMCMC: <0.001***  Age: 0.312(0.19 - 0.41)  pMCMC: <0.001*** | 1.49  (1.075 - 2) | 1.49  (1.07 - 1.97) |
| **1.2** | Intercept: -0.222(-0.35 - -0.063)  PMCMC: 0.0027***  Age: -0.011(-0.067 - 0.044)  pMCMC: 0.7 | Intercept: -0.222(-0.363 - -0.08)  pMCMC: 0.00533**  Age: -0.012(-0.071 - 0.043)  pMCMC: 0.68 | 1 | 1 |
| **2.1** | Intercept: -3.14 (-3.57- -2.72)  pMCMC: <0.001 ***  Age: 0.359 (0.24 - 0.48)  pMCMC: <0.001*** | Intercept: -3.14(-3.56 - -2.73)  pMCMC: <0.001  Age: 0.3562(0.23 - 0.47)  pMCMC: <0.001*** | 1.15  (0.8 - 1.6) | 1.14  (0.79- 1.62) |
| **2.2** | Intercept: -0.227(-0.361 - -0.059)  pMCMC: 0.0117***  Age: -0.01(-0.067 - - 0.0467)  pMCMC: 0.72 | Intercept: -0.226(-0.376 - -0.067)  pMCMC: 0.00333 ***  Age: -0.01(-0.066 --0.045)  pMCMC: 0.71 | 1 | 1 |
| **3.1** | Intercept: -3.16(-3.65 - -2.72)  pMCMC: <0.001***  Age: 0.369 (0.26- 0.51)  pMCMC: <0.001*** | Intercept: -3.18 (-3.7 - -2.7)  pMCMC: <0.001 ***  Age: 0.37 (0.25 - 0.5)  pMCMC: <0.001*** | 1.163 (0.74-1.58) | 1.158  (0.73 - 1.35) |
| **3.2** | Intercept: -0.22(-0.38 - -0.061)  pMCMC: 0.001**  Age: -0.01(-0.067 - 0.04)  pMCMC: 0.757 | Intercept: -0.226(-0.39 - -0.067)  pMCMC: 0.002***  Age: -0.009(-0.068- 0.046)  pMCMC: 0.776 | 1 | 1 |
| **4.1** | Intercept: -3.15(-3.62 - -2.74)  pMCMC: <0.001***  Age: 0.37 (0.256-0.49)  pMCMC: <0.001 *** | Intercept: -3.155(-3.6 - -2.7244)  pMCMC: <0.001***  Age: 0.37(0.25 - 0.48)  pMCMC: <0.001 *** | 1.159  (0.75 - 1.56) | 1.162  (0.76- 1.57) |
| **4.2** | Intercept: -0.23(-0.39 - - 0.067)  pMCMC: 0.005**  Age: -0.01(-0.068 - 0.045)  pMCMC: 0.728 | Intercept: -0.226(- 0.38 - -0.065)  pMCMC: 0.005**  Age: -0.01(-0.068 - 0.05)  pMCMC: 0.71 | 1 | 1 |
| **5** | Male Intercept: -3.24 (-3.7 - -2.84)  pMCMC: <0.001***  Male:Age: 0.328 (0.2 - 0.44)  pMCMC: <0.001**  Female Intercept: -0.217(-0.35 - -0.07)  pMCMC: <0.002**  Female:Age: -0.01(-0.07 - 0.04)  pMCMC: 0.64 | Male Intercept: -3.051(-3.45- -2.67)  pMCMC: <0.001***  Male Age: 0.31(0.2 - 0.42)  pMCMC: <0.001 ***  Female Intercept: -0.22(-0.365- -0.06)  pMCMC: 0.004**  Female Age: -0.012(-0.06 - 0.047)  pMCMC: 0.688 | Male: 1.952 (1.4 - 2.5)  Female: 1 | Male: 1.47  (1.075- 1.92)  Female: 1 |
| **6** | Male Intercept: -3.16(-3.57 - -2.73)  pMCMC: <0.001 ***  Male Age: 0.358 (0.23- 0.47)  pMCMC: <0.001 **  Female Intercept: -0.23(-0.39 - -0.08)  pMCMC: 0.005 ***  Female Age: 0.01(-0.07 - 0.04)  pMCMC: 0.64 | Male Intercept: -3.16(-3.56 - -2.71)  pMCMC: <0.001***  Male Age: 0.358(0.251 - 0.49)  pMCMC: <0.001***  Female Intercept: -0.226(-0.38 - -0.067)  pMCMC: 0.004 ****  Female Age: -0.01(-0.066 - 0.047)  pMCMC: 0.747 | Male: 1.165  (0.78 - 1.58)  Female: 1 | Male:  1.15  (0.71 - 1.55)  Female:  1 |
| **7** | Male Intercept: -3.17 (-3.65 - -2.67)  pMCMC: <0.001 ***  Male Age: 0.369 (0.243 - 0.48)  pMCMC: <0.001 ***  Female Intercept: -0.231 (-0.37 - -0.067)  pMCMC: <0.001  Female Age: -0.009 (-0.068 - 0.04)  pMCMC: 0.74 |  | Male:  Female: 1 | Male:  Female: 1 |

**Table S6: DIC, phenotypic scale heritability (H^2^_phen_), latent scale heritability (H^2^_lat_), phenotypic scale genetic correlations (Vg_phen_) and latent scale genetic correlations (Vg_lat_) from the posterior means of Prior 2 bivariate models. Heritability estimates where CIs were ≥0.001 are highlighted in bold. Residual and Fixed effect estimations can be found in Table S5.**

| **Models** | **Response(s)** | **Parameters** | **DIC** | **H^2^_phen_** | **H^2^_lat_** | **Vg_phen_** | **Vg_lat_** |
| --- | --- | --- | --- | --- | --- | --- | --- |
| **5** | Male EPR and female EPR | **cov(male DGE + female DGE) + idh(male PE + female PE)** | 4171 | Male: 0.004  (<0.001 - 0.015)  Female: 0.013  (<0.001 - 0.045) | Male: 0.026  (<0.001 - 0.094)  Female: 0.021  (<0.001 - 0.072) | 0.024  (-0.76 - 0.79) | 0.024 (-0.76 - 0.79) |
| **6** | Male EPR and female EPR | **cov(male DGE + female DGE) + idh(male PE + female PE) + idh(female partner PE + male partner PE)** | 4159 | Male: 0.006  (<0.001 - 0.012)  Female: 0.015  (<0.001 - 0.045) | Male: 0.041  (<0.001 - 0.14)  Female: 0.023 (<0.001 - 0.074) | 0.029 (-0.73 - 0.791) | 0.029 (-0.73 - 0.791) |
| **7** | Male EPR and female EPR | **cov(male DGE + female DGE) + idh(male PE + female PE) + idh(female partner PE + male partner PE) + cov(female partner IGE + male partner IGE)** |  | Male:  Female: | Male:  Female: |  |  |
